# Defects in nephrogenesis result in an expansion of the *Foxd1*+ stromal progenitor population

**DOI:** 10.1101/2025.02.10.637031

**Authors:** Michael G. Michalopulos, Yan Liu, Dinesh Ravindra Raju, John T. Lafin, Yanru Ma, Dhruv Gaur, Sadiksha Khadka, Chao Xing, Andrew P. McMahon, Thomas J. Carroll, Keri A. Drake

**Author notes:** **Corresponding author:** Keri Drake.

## Abstract

Reciprocal signaling interactions coordinate multiple aspects of kidney development. While signals from the stroma have been shown to regulate nephron progenitor cell (NPC) differentiation, much less is known about regulation of the stromal progenitor population. Here, we demonstrate that disruption of the NPC lineage via loss of *Wt1* (i.e., *Six2cre;Wt1^c/c^*) results in an expansion of *Foxd1*+ stromal progenitor cells. Analyses of the developing stroma in two additional models, including *Wnt4-null* mutants (which fail to form nephron structures similar to *Six2cre;Wt1^c/c^* kidneys) and NPC ablation via diphtheria toxin (i.e., *Six2cre;RosaDTA^c/+^*), both phenocopy *Six2cre;Wt1^c/c^* mutants, thus further confirming that defects in the NPC lineage result in abnormal development of the stromal progenitor population. Furthermore, we identify a subcluster of the *Foxd1*+ stroma that appears expanded in the three mutant mouse models and conserved in human fetal kidneys. Overall, the findings from this study suggest that loss of differentiating nephron structures may result in possible over proliferation of the stromal progenitor population and/or a block in stromal differentiation and further highlight how crosstalk amongst the progenitor cell lineages coordinates multiple aspects of kidney development.

**KEY POINTS:**

- Mutant mouse models targeting the nephron lineage (i.e., *Six2cre;Wt1^c/c^* and *Wnt4-null* mutants which fail to form early nephron structures) suggest that a block in NPC differentiation results in an abnormal expansion of the *Foxd1*+ stromal progenitor population.
- Mutant kidneys with NPC ablation (i.e., *Six2cre;RosaDTA^c/+^*) show maintenance of stromal progenitors independent of signals from adjacent nephron progenitors and ureteric bud.
- Single nuclei RNA-seq identifies three subclusters of the *Foxd1*+ stromal progenitor population at E15.5, including one cluster of proliferating cells and a distinct *Fap*+, *Igf1*+, *Gria1*+, *Gdnf*– subcluster which appears expanded in *Six2cre;Wt1^c/c^*, *Wnt4-null,* and *Six2cre;RosaDTA^c/+^* mutant kidneys and conserved in human fetal kidneys.

## INTRODUCTION

The development of the mammalian kidney is a highly coordinated process involving reciprocal signaling interactions among the nephron, ureteric, and stromal progenitor lineages. Such cell-lineage crosstalk has been shown to regulate multiple aspects of kidney development, from directing the initial events in the formation of the metanephros via invasion of the ureteric bud (UB) into the metanephric mesenchyme (Grobstein, 1953, 1956) to regulating the self-renewal, proliferation, and differentiation of nephron progenitor cells (NPCs) and UB branching essential in determining final nephron allotment (Barak et al., 2012; Carroll et al., 2005; Costantini & Kopan, 2010). However, the extent to which cellular crosstalk coordinates the complex processes in normal kidney development is still being uncovered.

The self-renewing *Foxd1*+ stromal progenitor population gives rise to the majority of the renal interstitium, including pericytes, mesangial cells, and vascular smooth muscle (Kobayashi et al., 2014). Signals from the embryonic stroma have been shown to regulate NPC maintenance/differentiation, UB branching, and patterning of the vasculature via non-cell autonomous mechanisms (Das et al., 2013; Hatini et al., 1996; Hum et al., 2014; Levinson et al., 2005). Additionally, the integration of lineage-specific stromal cells in kidney organoids has further confirmed the importance of the stroma in generating “higher-order structures” recapitulating in-vivo development (Tanigawa et al., 2022). Despite this exciting progress, much remains to be understood about mechanisms regulating stromal progenitor maintenance and differentiation as well as the potential contributions to diseases that may be associated with stromal defects in both developing and adult kidneys.

While NPC differentiation is known to be regulated by signals from the stroma, much less is known about mechanisms regulating the maintenance/differentiation of the stromal progenitor population. Here we sought to further examine how defects in the nephron lineage may non-autonomously affect stromal development using in-vivo mouse models targeting mutations to the NPCs. Specifically, we examined three genetically engineered mouse lines including 1) *Six2cre;Wt1^c/c^* mutant kidneys, given that loss of *Wt1* in the NPCs has been shown to block NPC differentiation with maintained self-renewing NPCs, 2) *Wnt4-null* mutants, which also fail to undergo NPC differentiation/mesenchyme-to-epithelial (MET) transition but avoids targeting of the self-renewing NPC population, and 3) NPC ablation via diphtheria toxin using the *Six2cre;RosaDTA^c/+^* model. Evaluation of stromal markers in these mouse models demonstrates an increase of the nephrogenic zone stroma by immunofluorescence (IF) and in-situ hybridization (ISH) of known stromal progenitor markers including forkhead box D1 (*Foxd1*) and netrin 1 (*Ntn1*), as well as newly identified markers from E15.5 single nuclei RNA-seq (snRNA-seq) including early B cell factor 3 (*Ebf3*), aldehyde oxidase 3 (*Aox3*), fibroblast activation protein (*Fap*), insulin growth factor 1 (*Igf1*), and glutamate ionotropic receptor AMPA type subunit 1 (*Gria1*). We also show the expansion of a distinct *Fap*+, *Igf1*+, *Gria1*+, *Gdnf*– subcluster of the *Foxd1*+ stromal progenitor population, which is maintained independent of signals from the NPCs/UB as shown in the *Six2cre;RosaDTA^c/+^* model and appears to be conserved as subcluster of stromal cells in human fetal kidneys. While the abnormal expansion of the stromal progenitor population does not appear to be solely due to a “delay” in development, further studies will be needed to understand if this expansion potentially results from unrestrained proliferation vs a possible block in stromal progenitor differention. Nonetheless, the findings from this study suggest coordination in the development of nephron and stromal lineages, with defects in NPC differentiation resulting in non-cell autonomous effects on the stromal progenitor population, and additionally identify a novel subcluster of the *Foxd1*+ stromal progenitor population that appears to be conserved in the human fetal kidney.

## RESULTS

### Genetically engineered mouse models disrupting the NPC lineage show an abnormal expansion of the nephrogenic zone stroma

Previous studies have shown that loss of *Wt1* in the nephron lineage prevents nephrogenesis, with failed MET through the transcriptional regulation of *Wnt4* (Berry et al., 2015; Davies et al., 2004; Essafi et al., 2011). Here, we targeted a conditional *Wt1* mutation with *loxP* sites flanking exons 8 and 9 (Gao et al., 2006) to the NPCs using *Six2cre* (Kobayashi et al., 2008). We confirmed that *Six2cre;Wt1^c/c^* mutant kidneys maintain SIX2+ NPCs (Fig. 1, panel F and Supplemental Fig. S1) and show defects in NPC differentiation (Fig. 1, panel J, with NCAM+ early differentiating structures marked by arrows, and Supplemental Fig. S1, panel F showing loss of LHX1 structures marked by arrowheads). While *Six2cre;Wt1^c/c^* mutant kidneys do form some differentiating structures with observed proximal tubules and glomeruli at later timepoints (Supplemental Fig. S1, panel B, arrows), this is presumably due to incomplete efficiency of the *Six2cre* model which has been previously reported with some NPCs “escaping” cre-recombination (Marable et al., 2018) and/or incomplete *loxP* site excision targeting two alleles in this homozygous mutant model. Inclusion of a *Rosa26^EYFP^* reporter was used to confirm that *Six2cre* specifically targets the NPC lineage with no recombination observed in the stroma as expected (Supplemental Fig. S1, panel H). We next examined H&E images of *Six2cre;Wt1^c/c^* mutant kidneys, which suggest an expansion of cells in the nephrogenic zone of mutant kidneys surrounding the cap mesenchyme (Fig. 1, panel B). IF of the pan-stromal marker meis homeobox paralogs 1, 2, 3 (MEIS1/2/3) and the nephrogenic stromal marker aldehyde dehydrogenase 1 family member A2 (ALDH1A2) indeed shows an expansion of stroma, appearing as a “multi-layering” of stromal cells, between the outer cap mesenchyme and periphery of the kidney (Fig. 1, panels F and N, respectively), where the *Foxd1*+ stromal progenitor population normally resides. Additionally, NCAM expression appears to be ectopically expressed in this nephrogenic zone stroma of *Six2cre;Wt1^c/c^* mutant kidneys (Fig. 1, panel J, arrowhead).

**Figure 1.**
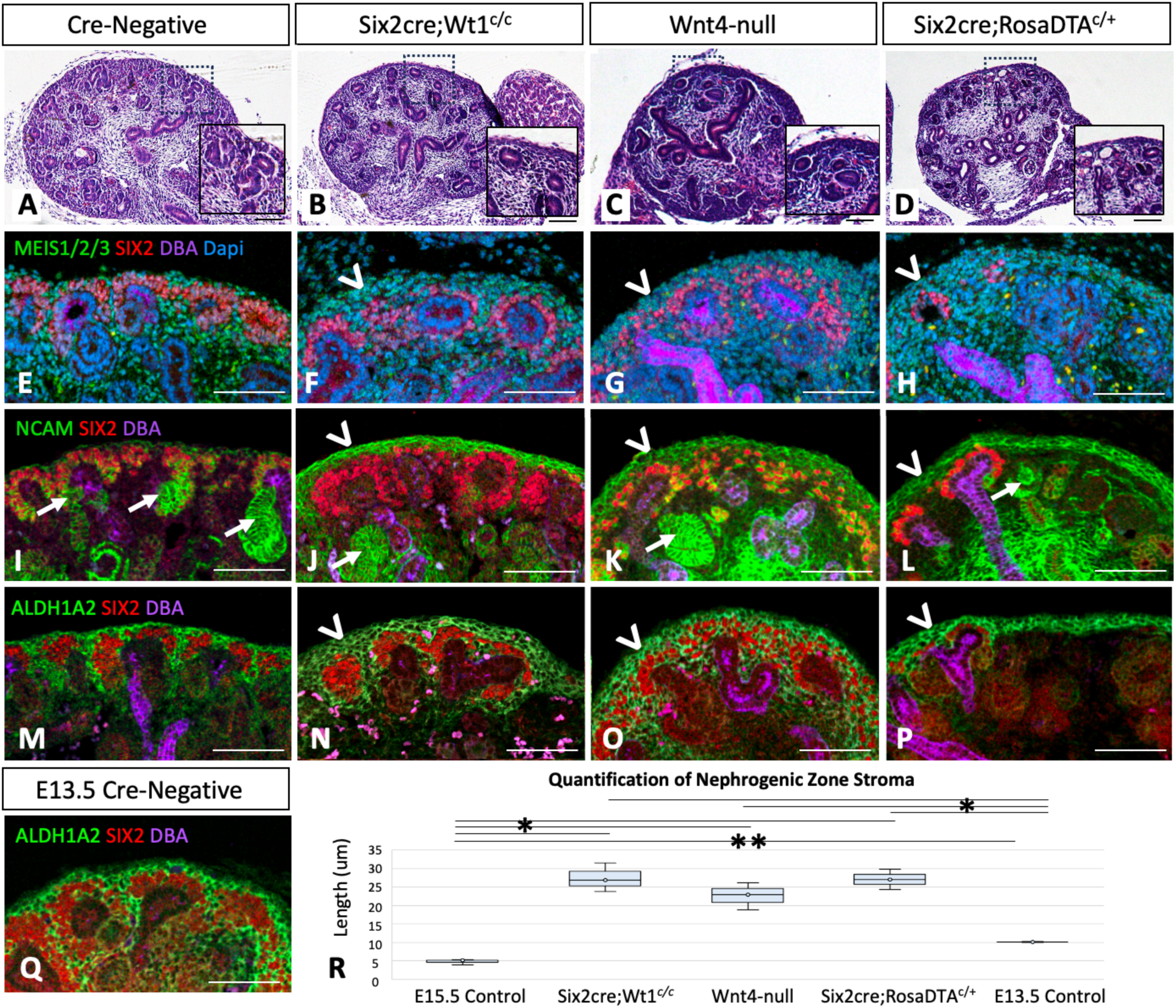
Mutant mouse models with defects in nephrogenesis show an expansion of stroma in the nephrogenic zone. (A-D) Histology of E15.5 cre-negative control kidneys (A) is shown in comparison to mutants models with defects in nephrogenesis including *Six2cre;Wt1^c/c^* (B), *Wnt4-null* (C) and *Six2cre;DTA^c/+^* (D); scale bars = 100 um; representative H&E images shown from N=3. (E-Q) Immunofluorescence of E15.5 control and mutant kidneys show NPCS (labed with SIX2), early differentiating nephron structures (labeled with NCAM, arrows), ureteric bud/collecting ducts (labeled with DBA), stroma (with MEIS1/2/3 as a global stromal marker of the stroma and ALDH1A2 specifically expressed in the stroma surrounding the cap mesenchyme, or “nephrogenic zone”). Mutant kidneys with defects in nephrogenesis show expansion of stromal cells at the outer periphery of the kidney (arrowheads) with abnormal expression of NCAM (J-L) and expression ALDH1A2, a marker of the nephrogenic zone stroma (N-P). ALDH1A2 expression at earlier timepoints in development (E13.5 shown here) does not show similar expansion (Q); scale bars = 50 um; representative images shown from N=3. (R) Quantification of the nephrogenic zone stroma was performed by using image J to measure the distance from the outer cap mesenchyme to the periphery of the kidney on sections of control and mutant kidneys stained with ALDH1A2 and SIX2 (i.e., as shown in M-P); * p-value < 0.001, ** p-value = 0.03.

Given that loss of *Wt1* in the nephron lineage results in a block in NPC differentiation as well as effects on gene expression in the self-renewing NPCs (Berry et al., 2015), we sought to examine additional mutant models disrupting the nephron lineage to evaluate whether effects on the nephrogenic zone stroma are specific to *Six2cre;Wt1^c/c^* mutant kidneys or may be seen in other developmental models that disrupt the nephrogenesis. To do this, we examined two additional models, including *Wnt4-null* and *Six2cre;RosaDTA^c/+^* transgenic mice. Specifically, loss of *Wnt4* is known to disrupt NPC differentiation/tubulogenesis (Kispert et al., 1998; Stark et al., 1994), similar to mutant kidneys with loss of *Wt1* targeted to the nephron lineage but without the effects of *Wt1* mutation in the self-renewing NPCs. The *Six2cre;DTA^c/+^* transgenic mouse model results in the ablation of SIX2+ NPCs via apoptosis triggered by the cre-inducible DTA, diphtheria toxin fragment A (Brockschnieder et al., 2004). As expected, *Six2cre;DTA^c/+^* kidneys show severe defects in development including markedly decreased UB branching and very few differentiated tubules with sparse, residual NPCs (Fig. 1, D) presumed due to incomplete efficiency of the *Six2cre* model (Marable et al., 2018) as described above. Both models show expansion of the stroma by MEIS1/2/3 and ALDH1A2 expression at the outer periphery of the kidney (Fig. 1, panels G, H and O, P respectively) and also show ectopic expression of NCAM in the nephrogenic zone stroma (Fig. 1, panels K and L). Thus, this expansion and abnormal patterning of the developing stroma in *Wnt4-null* and *Six2cre;RosaDTA^c/+^* mutant kidneys appear to phenocopy the stromal changes identified in the *Six2cre;Wt1^c/c^* model.

Given that all three models cause severe disruptions to the developing kidney, we additionally evaluated ALDH1A2 expression at earlier timepoints in development to see if the expansion of nephrogenic zone stroma may be related to a “developmental delay” or an “immature” phenotype of the developing stroma. At E13.5 (Fig. 1, panel Q), there does not appear to be a similar expansion ALDH1A2 expression compared to E15.5 in the three mutant models. To further confirm this, we quantified the expansion of the nephrogenic zone stroma using image J to measure the distance from the outer cap mesenchyme (i.e., SIX2 posivite cells) to the periphery of the kidney on sections of control and mutant kidneys stained with ALDH1A2 and SIX2 (with representative images shown in Fig. 1, panels M-Q). Ten measurements were obtained from three separate embryos of each of the five sample types analyzed, with measurements from *Six2cre;RosaDTA^c/+^* mutants including both regions where the NPCs were present and regions in which they were completely ablated. From this analysis, *Six2cre;Wt1^c/c^*, *Wnt4-null,* and *Six2cre;RosaDTA^c/+^* mutant kidneys demonstrate a statistically significant increase in nephrogenic zone stromal length compared to both E15.5 and E13.5 control kidneys (p-value < 0.001). Additionally, E13.5 samples were also found to have increase length compared to E15.5 (p-value = 0.03); however, the average measurement from E13.5 kidneys was 10.1 μm, which is half that of the measurements from the three mutant models (with the average of *Six2cre;Wt1^c/c^*, *Wnt4-null,* and *Six2cre;RosaDTA^c/+^* mutants being 27.4, 22.6, and 27.1 μm respectively). No statistically significant differences were noted among the three mutant models.

### Single nuclei RNA-sequencing identifies subclusters of the *Foxd1*+ stromal progenitor population

As described above, the *Foxd1*+ stromal progenitor population resides specifically in the nephrogenic zone of the developing kidney where cellular expansion occurs in the *Six2cre;Wt1^c/c^*, *Wnt4-null,* and *Six2cre;RosaDTA^c/+^* mutant models. Given that previous studies profiling the *Foxd1*+ stromal progenitor cells have identified cellular heterogeneity within this progenitor population at E18.5 (England, 2020), we sought to evaluate snRNA-seq from E15.5 control kidneys to find additional markers of the *Foxd1*+ stromal progenitor cells at this earlier time point and include kidneys from *Six2cre* embyros as control samples for the *Six2cre;Wt1^c/c^* model. SnRNA-seq from three separate samples (i.e., paired kidneys from one cre-negative embryo and two *Six2cre* embryos) were pooled to total 15,636 nuclei after quality control filtering (Fig. 2, A). Cell type-specific markers (Fig. 2, B and Supplemental File S1) were utilized to label cell types based on marker gene lists and anchor gene expression (Chaney et al., 2022) after unsupervised cell clustering was performed, which identified a total of 18 clusters corresponding to specific cell types. Four “unidentified” clusters showed either relatively lower reads, high ribosomal expression, and/or lack of specific marker expression likely due to technical artifact and were not included in further analyses.

**Figure 2.**
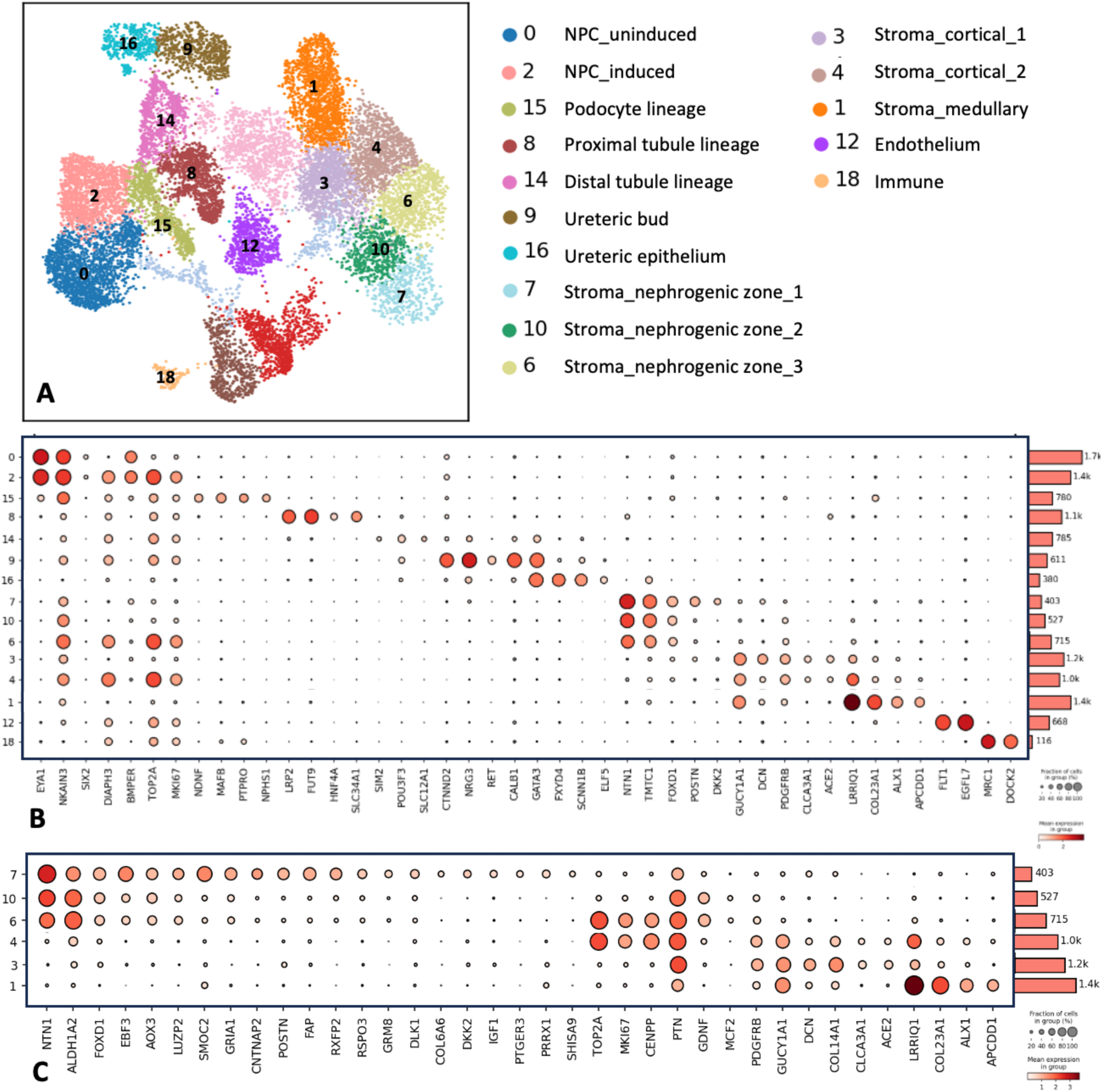
Single nuclei RNA-seq of control kidneys (cre-negative and *Six2cre*) at E15.5 identifies heterogeneity of the *Foxd1*+ stromal progenitor population. (A) A total of 15,636 nuclei from cre-negative kidneys (N=1) and *Six2cre* only kidneys (N=2) were pooled for analysis, with clustering of the nuclei shown via UMAP. (B) Cell type-specific markers (with representative genes shown in this dotplot) were used to identify 18 clusters/cell types and four “unidentified” clusters that showed low reads, high ribosomal expression, and/or lack of specific marker expression likely due to technical artifact and thus not included in further analyses. (C) Stromal clusters were specifically evaluated for gene expression profiles distinguishing the six clusters, with three clusters (7, 10, and 6) showing enriched *Foxd1*+ expression consistent with the stromal progenitor population.

Next, we specifically evaluated the six stromal clusters identified (totaling 5,216 nuclei) for distinguishing gene expression profiles, with three clusters (7, 10, and 6) showing enriched *Foxd1*+ expression consistent with the stromal progenitor population (totaling 1,645 nuclei). Minimal gene expression differences were identified between the stromal progenitor cells of the cre-negative kidneys vs the *Six2cre* kidneys (Supplemental File S2), so captured nuclei from all three control samples were pooled for subsequent analyses. The three *Foxd1*+ subclusters (clusters 7, 10, and 6) show enriched expression *of Ntn1* and *Aldh1a2* as expected, as well as additionally identified markers *Ebf3, Aox3, and Luzp2* (Fig. 2, panel C). Differentially expressed genes (DEGs) among the subclusters shows cluster 7 with enriched expression of *Smoc2, Gria1, Cntnap2, Postn, Fap, Rxfp2, Rspo3, Grm8, Dlk1, Dkk2, Igf1, Ptger3, Prrx1,* and *Shisa9.* Cluster 10 shows decreased expression of the above markers with upregulation of *Ptn, Gdnf, and Mcf2*, while cluster 6 shows enriched expression of proliferation markers including *Top2a, Mki67, and Cenpp* (Fig. 2, panel C and Supplemental File S3). The additional stromal clusters 4, 3, and 1 show gene expression profiles consistent with stroma localized to the cortical and medullary regions of the kidney, as marked by expression of *Gucy1a1, Clca3a1*, and *Ace2* in cortical stromal clusters 4 and 3, and *Lrriq1, Col23a1, Alx1*, and *Apcdd1* in medullary stromal cluster 1. Thus, this analysis identifies additional markers specific to the *Foxd1*+ stromal progenitor population and its subclusters that we could further evaluate in our mutant mouse models.

### The *Foxd1*+ stromal progenitor population, including the *Fap*+, *Igf1*+, *Gria1*+, *Gdnf*– subcluster identified from snRNA-seq, is abnormally expanded in mutant kidneys with defective nephrogenesis

We next examined the expression of markers specific to the *Foxd1*+ stromal progenitor population by ISH in *Six2cre;Wt1^c/c^*, *Wnt4-null,* and *Six2cre;RosaDTA^c/+^* mutant kidneys compared to cre-negative control kidneys (Fig. 3), including the known markers *Foxd1* and *Ntn1*, as well as the newly identified markers *Ebf3* and *Aox3*. Markers of subcluster 7 including *Fap*, *Igf1*, and *Gria1* and subclusters 6 and 10 including *Gdnf* and *Ptn* were also examined. Indeed, *Foxd1*, *Ntn1*, *Ebf3* and *Aox3* were expanded in *Six2cre;Wt1^c/c^*, *Wnt4-null,* and *Six2cre;RosaDTA^c/+^* mutant kidneys. The outer periphery of all three mutant kidney models also show increased expression of *Fap*, *Igf1*, and *Gria1* stromal cells that lack expression of *Gdnf* and *Ptn*, consistent with the snRNA-seq gene expression profile of subcluster 7 of the *Foxd1*+ stromal progenitor population.

**Figure 3.**
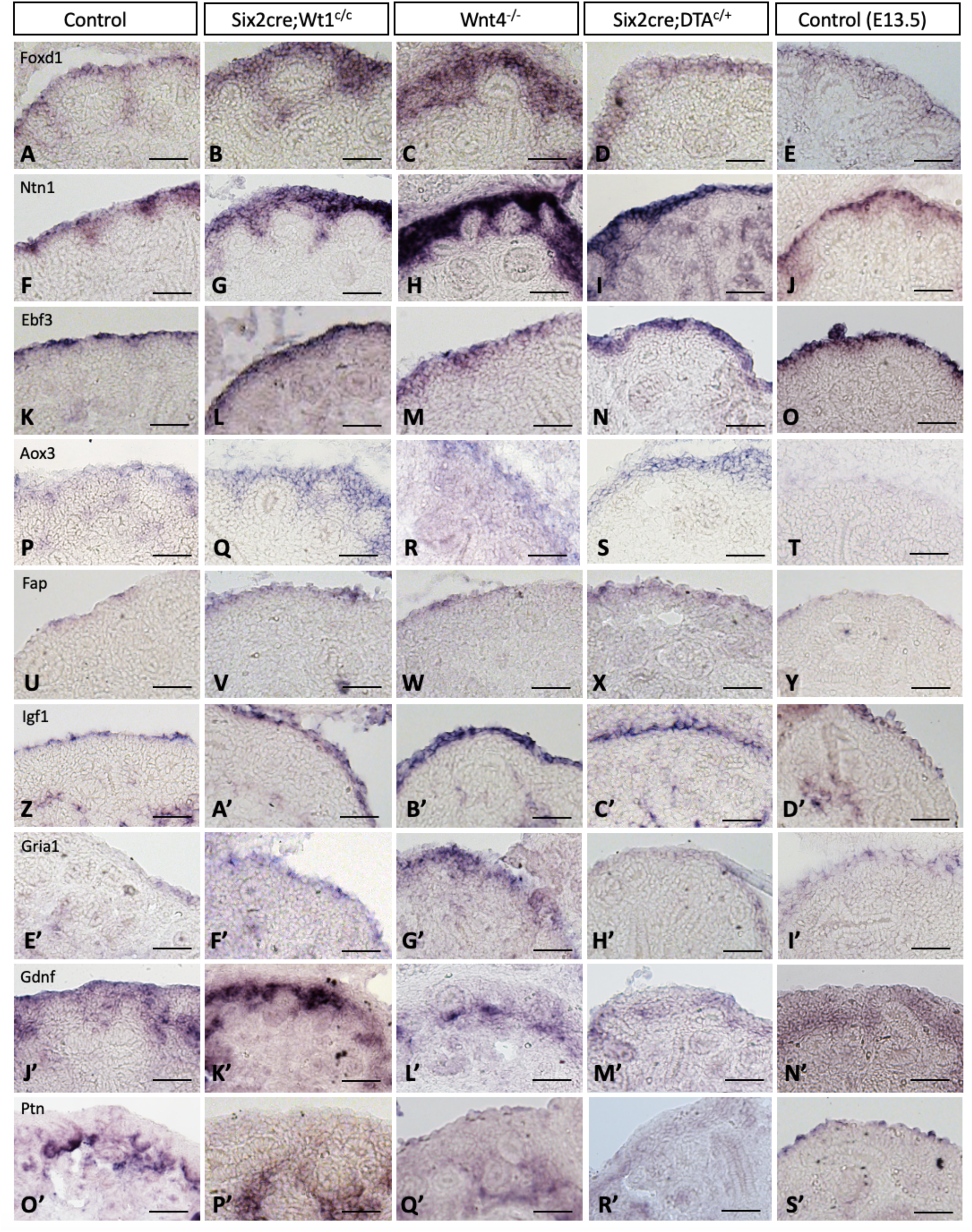
The *Foxd1+* stromal progenitor population shows abnormal patterning in mutant kidneys with defects in nephrogenesis. In-situ hybridization of known markers of the nephrogenic zone stroma, *Foxd1* and *Ntn1*, as well as markers identified from snRNA-seq analyses including *Ebf3, Aox3, Fap, Igf1,* and *Gria1*, all show expanded/increased expression in *Six2cre;Wt1^c/c^*, *Wnt4-null*, and *Six2cre;RosaDTA^c/+^* mutant compared to control kidneys at E15.5. Localization of *Gdnf* expression shows minimal signal in the outer periphery of the nephrogenic stroma of mutant kidneys vs E15.5 controls, with a similar expression pattern of *Ptn*, suggesting an expansion of subcluster 7 of the *Foxd1*+ stroma identified from our snRNA-seq dataset showing *Fap*+, *Igf1*+, *Gria1*+, *Gdnf*–. When comparing the expression of these markers to E13.5 control kidneys, *Foxd1* and *Ntn1* appear abnormally expanded in mutant kidneys vs E13.5 controls, with additional control images of early timepoints shown in Supplemental Figure S1. However, *Fap*, *Igf1*, *Gria1*, and *Gdnf* appear similar to mutant kidneys, suggesting that subcluster 7 may be expanded at earlier timepoints development, yet the restrticted expression of *Foxd1* and *Ntn1* at E13.5 indicates abnormal patterning of stromal progenitor cells in E15.5 mutant kidneys. Scale bars = 50 um; representative images shown from N=3.

We additionally examined the expression of these markers at earlier time points in development to further evaluate if the changes in stromal expression may reflect an “immature” pattern of the developing stroma, as described above. Notably, *Foxd1*, *Ntn1*, and *Aox3* do not appear expanded at E13.5 in comparison to control kidneys at E15.5 (Fig. 3, panels E, J, and T respectively). However, *Ebf3*, *Fap*, and *Gria1* do appear somewhat expanded in control kidneys at E13.5 in comparison to more restricted expression at E15.5 (Fig. 3, panels O, Y, and I’ respectively). Furthermore, *Gdnf* shows lower levels of expression and appears excluded from the outer peripheral cells of the nephrogenic zone stroma at E13.5 (Fig. 3, panel N’). Given these developmental-specific expression patterns, we systematically evaluated the localization of *Foxd1, Ntn1, Ebf3, Aox3, Fap, Igf1, Gria1, Gdnf*, and *Ptn* throughout development at E12.5, E13.5, E15.5, and E18.5 (Supplemental Fig. S2), with some temporal-specific expression patterns observed. Taken together, these findings suggest that the expansion of subcluster 7 of *Foxd1*+ progenitor cells in mutant kidneys may reflect an “immature” phenotype of the stromal progenitor population; however, the significant expansion of *Foxd1*+ and *Ntn1*+ cells in mutant kidneys (that is not seen at earlier time points E12.5 and E13.5) suggests that defects in nephrogenesis additionally result in an abnormal expansion of the stromal progenitor population not soley explained by a “delay” in developmental maturation.

Since the *Six2cre;RosaDTA^c/+^* model results in incomplete NPC ablation as described above, we carefully examined the expression of markers of the stromal population in both regions of complete NPC ablation as well as regions where residual NPCs were present (Fig. 4, panels B and D). Interestingly, *Six2cre;DTA^c/+^* mutant kidneys show stromal progenitor cells maintained throughout the periphery of the kidney even in regions of complete NPC ablation, as suggested by expression of ALDH1A2 (Fig. 4, panel D, arrowheads), with no significant difference in the quantification of nephrogenic zone stromal length in regions of retained NPCs vs complete NPC ablation as quantified by image J described above (Fig. 4, panel E). Additionally, ISH shows expression of *Foxd1, Ntn1, Fap,* and *Igf1* with a lack of *Gdnf* expression (Fig. 4) in the outer periphery stroma of these mutant kidneys, suggesting that cells from subcluster 7 of the *Foxd1*+ stromal progenitor population are maintained in *Six2cre;DTA^c/+^* mutant kidneys, even in regions with complete loss of the NPCs/UB of the nephrogenic niche.

**Figure 4.**
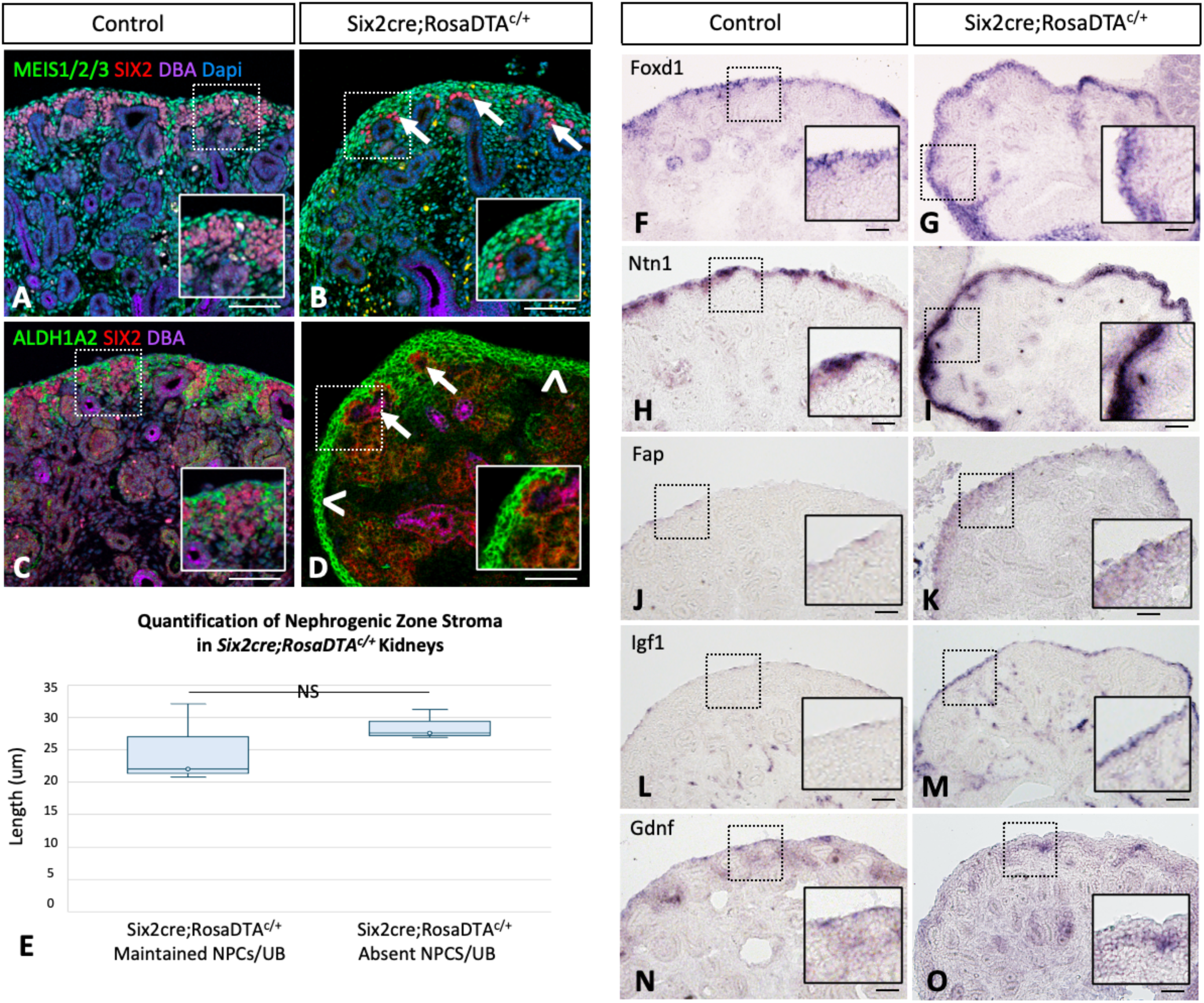
Stromal progenitor cells are maintained independent of the nephrogenic niche in *Six2cre;RosaDTA^c/+^* mutant kidneys. (A-D) In comparison to control kidneys, *Six2cre;RosaDTA^c/+^* mutant kidneys show expanded stroma throughout the periphery of the kidney as described above. Examination of both regions where the NPCs/UB persist due to incomplete NPC ablation with the *Six2cre* model (arrows) and in regions of complete ablation (arrowheads) confirm expansion. (E) Quantification of the nephrogenic zone stroma (as described in Figure 1) in both regions in which NPCs/UB are maintained vs regions of their absence did not show a statistically significant difference (NS; p-value 0.49). (F-O) In-situ hybridization of *Foxd1, Ntn1, Fap*, *Igf1*, and *Gdnf* additionally show that the maintained stromal progenitor cells express markers of subcluster 7 identified in the snRNA-seq dataset showing heterogeneity of the *Foxd1*+ stroma progenitor population. Scale bars = 50 um; representative images shown from N=3.

### Human fetal kidneys show conserved expression of several murine stromal progenitor cell markers

While comparative studies have shown many conserved features between mouse and human kidney development, recent studies have identified distinct gene expression profiles suggesting important divergent features as well (Kim et al., 2024; Lindström, McMahon, et al., 2018; Lindström, Tran, et al., 2018). To evaluate if the heterogeneity identified in the murine *Foxd1*+ stromal progenitor population may be conserved in human fetal kidney development, we re-analyzed publically available human fetal kidney snRNA-seq data (Kim et al., 2024) to evaluate for the expression of our identified murine stromal progenitor cell markers. To do this, we first performed unsupervised cell clustering and labeled cell types based on marker gene lists and anchor gene expression, or informed cell clustering (Chaney et al., 2022). Within the human fetal snRNA-seq dataset (Fig. 5, panels A and B, and Supplemental File S4), we identified six stromal clusters (i.e., clusters 1, 2, 6, 11, 12, and 23) totaling 11,744 nuclei. Using the known markers identified in murine (England, 2020) and human kidneys (Kim et al., 2024), two clusters, including 1 and 11, show enriched expression of the nephrogenic zone stromal markers *NTN1* and *FOXD1*, while cluster 12 shows increased expression of proliferation markers including *TOP2A* and *MIK67*, and clusters 2, 23, and 6 show enriched expression of cortical and medullary stromal markers *GUCY1A1*, *COL23A1*, *APCDD1*, and *ALX1*. Further examination of the *NTN1*+, *FOXD1*+ stromal cells identifies specific expression of *COL14A1* and *MGAT4C* in cluster 1 and *MYOCD* and *FST* showing specific expression in cluster 11 (Fig. 5, panel B).

**Figure 5.**
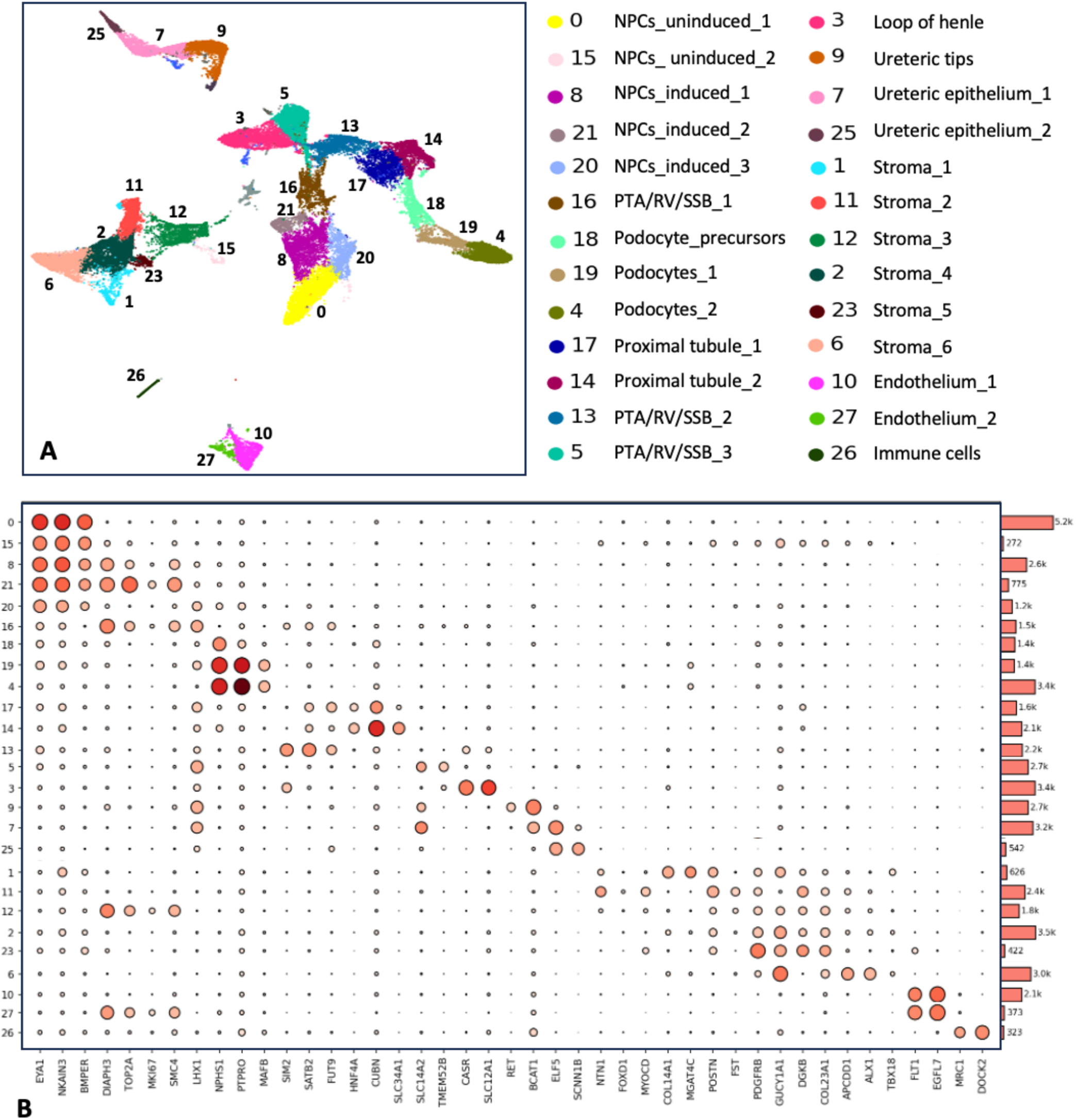
Human fetal kidney snRNA-seq identifies distinct stromal cell types. (A) Publically available snRNA-seq including 52,046 nuclei from human fetal kidneys was re-analyzed, as shown in this UMAP, to specifically identify subclusters of human fetal kidney stroma. (B) Cell type-specific markers (with representative genes shown in this dotplot) were used to label cell types after performing unsupervised clustering, which identified a total of 27 clusters/cell types, including six clusters of stroma (i.e., clusters 1, 2, 6, 11, 12, and 23) showing distinct expression profiles.

Next, we further evaluated the transcriptional profiles of the 3,027 nuclei from the *NTN1* and *FOXD1* expressing clusters 1 and 11, specifically evaluating the expression of the murine stromal progenitor cell markers identified from our snRNA-seq analysis (Fig. 6). Indeed, cluster 7 of the murine snRNA-seq dataset and cluster 1 of the human fetal kidney snRNA-seq dataset both show enriched expression of a number of markers, including *EBF3, LUZP2, COL6A6, PRRX1, IGF1, DKK2, RXFP2, FAP, RSPO3, SMOC2*, and *POSTN* with low *GDNF* expression. However, other murine markers, including *Aldh1a2*, *Cntnap2, Gria1, Dlk1, Grm8, Ptger3, Shisa9*, and *Aox3* show divergent expression patterns between the murine and human datasets. To visualize the conserved vs divergent patterns in expression of these markers, we evaluated the expression of specific markers genes across all the stromal clusters (Fig. 6), with conserved genes highlighted in green, divergent genes highlighted in red, and murine-specfiic genes highlighted in gray. Additional marker genes analysis (Supplemental File S1 and S4) and heatmap visualization (Supplemental Fig. S3) suggest distinct transcriptional profiles among the identified murine and human stromal clusters, consistent with previous mouse studies demonstrating heterogeneity of the interstitial cell types at later embryonic time points (England, 2020). While additional studies are needed to further validate the expression of stromal marker genes in human fetal kidneys, this snRNA-seq analysis identifies a number of novel markers that appear specific to the stromal progenitor cells of the developing kidney in both humans and mice and may offer the opportunity to better understand the molecular signature and unique functions of this specialized progenitor population.

**Figure 6.**
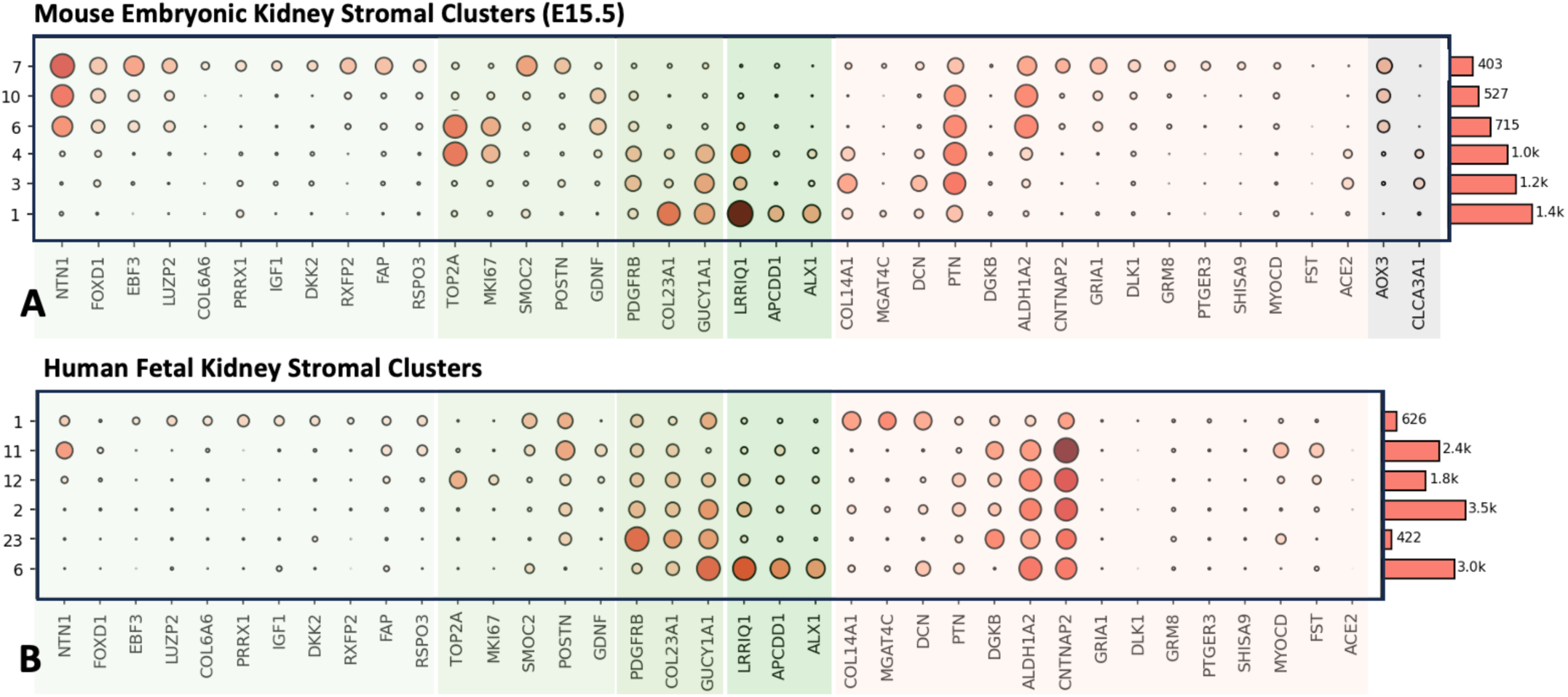
While comparisons of mouse and human stromal cell types identified via snRNA-seq suggests both conserved and divergent gene expression between species, these findings provide insights into the transcriptional profiles of the stromal progenitor population in both humans and mice. (A) As previously described, snRNA-seq identifies distinct transcriptional profiles for the stromal progenitor population (clusters 1, 10, and 6), cortical stroma (clusters 4 and 3), and medullary stroma (cluster 1). When comparing the transcriptional markers of these stromal cell clusters to the cell clusters identified in the human fetal kidney data set (B), some markers show similar expression profiles between the species (highlighted in green), whereas other markers do not show conserved expression (highlighted in red), and two markers were only identified in mouse stroma and not expressed in humans (highlighted in gray). Notably, stromal subcluster 1 in the human dataset shows expression of *Ntn1*, *Fap*, *Igf1*, as well as a number of other markers (including low *Gdnf* expression) similar to subcluster 7 of the *Foxd1+* stromal progenitor population in mice.

## DISCUSSION

While it has long been recognized that interactions among the NPCs, stromal progenitor cells, and UB play critical roles in regulating kidney development, the extent to which such cell-lineage crosstalk coordinates the intricate events of nephrogenesis is still being understood. Here, we show that mouse models with defects in nephrogenesis (i.e., *Six2cre;Wt1^c/c^*, *Wnt4-null*, and *Six2cre;RosaDTA^c/+^*) demonstrate an abnormal expansion of the *Foxd1*+ stroma, including cells from a distinct subcluster of this progenitor population (as summarized in Fig. 7) that appears to be conserved in human fetal kidney development.

**Figure 7.**
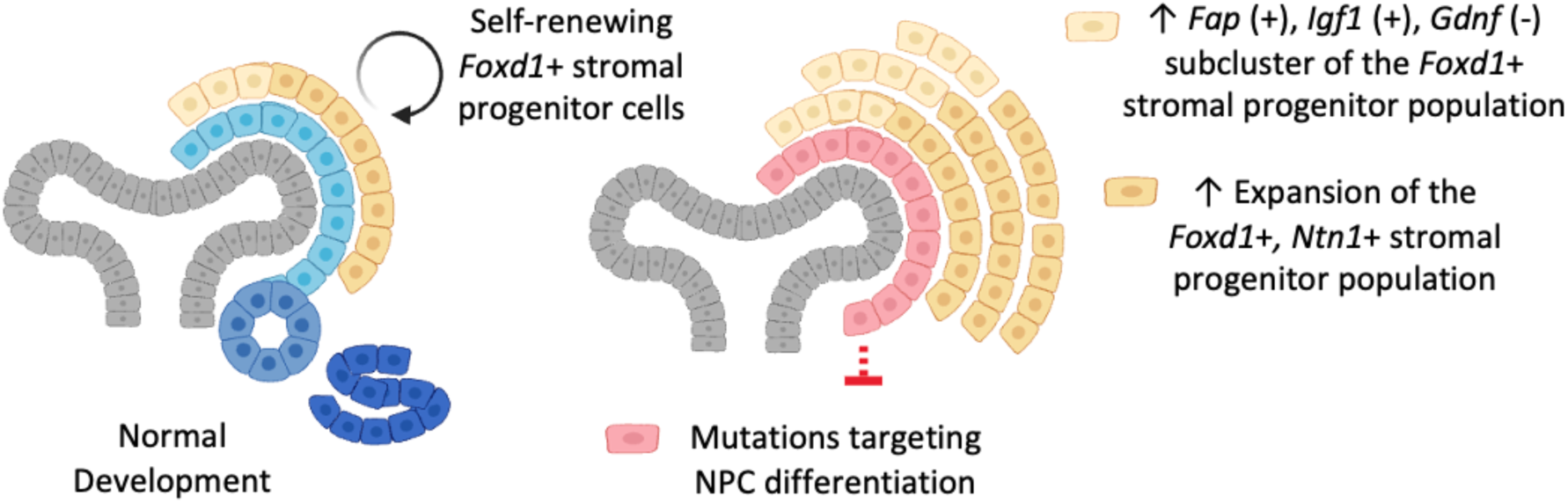
Model of stromal progenitor expansion in mutant kidneys with defects in nephrogenesis. Findings from three mouse mutant models (i.e., *Six2cre;Wt1^c/c^*, *Wnt4-null, Six2cre;RosaDTA^c/+^*) suggest that disruptions in nephron progenitor differentiation result in an abnormal expansion of the *Foxd1*+ stromal progenitor population, including a distinct *Fap* (+), *Igf1* (+), *Gdnf* (-) subcluster that appears to be conserved in human fetal kidneys and may further characterize the stromal progenitor population of both the human and mouse kidney.

### The *Foxd1*+ stromal progenitor population is heterogeneous, with a specific subset of cells (showing a distinct transcriptomic profile) maintained independent of signals from the nephrogenic niche

Heterogeneity of the *Foxd1*+ stromal progenitor population has been previously suggested via single-cell RNA-sequencing of murine kidneys at E18.5 (England, 2020). Similarly, our snRNA-seq analysis of the *Foxd1*+ stroma from control kidneys at E15.5 identifies three subclusters, including a distinct subcluster localized to the outer periphery of the nephrogenic zone stroma marked by expression of *Fap, Igf1, Gria1, Cntnap2, Postn, Rxfp2, Rspo3, Grm8, Col6a6*, and *Shisa9,* with decreased expression of *Gdnf* and *Ptn*. Interestingly, this specific subset of the *Foxd1*+ stromal progenitor population appears expanded in *Six2cre;Wt1^c/c^*, *Wnt4-null*, and *Six2cre;RosaDTA^c/+^* mutant kidneys (Fig. 3) and is maintained independent of signals from the NPCs/UB in the nephrogenic niche (Fig. 4). This is also similar to what occurs to the NPCs in the stromal-ablation model, as apoptosis of the stromal progenitor cells with diphtheria toxin (i.e., *Foxd1cre;RosaDTA^c/+^* mutant kidneys) results in maintained self-renewing NPCs that fail to differentiate and actually show an “expansion” of their population due to the loss of signals from the developing stroma (Das et al., 2013; Hum et al., 2014). These findings thus suggest that neither the nephron progenitor nor the stromal progenitor populations rely on signals from one another for their survival. While further studies will be necessary to evaluate if the abnormal expansion of the stromal progenitor population may occur due to unrestrained proliferation vs a possible block in stromal progenitor differention, the findings from this study provide an additional example of coordination in the development of nephron and stromal lineages. Additionally, validation of specific marker genes via in-situ hybridization along with the transcriptomic profiling by snRNA-seq raise the possibility that we have identified a distinct subset of the multipotent progenitor cells within the *Foxd1*+ stroma, with a number of specific markers of this *Foxd1*+ subcluster appearing conserved in the human fetal kidney. While outside the scope of this report, future studies will be critical to further evaluate this finding and may offer the opportunity to develop novel tools and better understand mechanisms governing the development of the renal stroma.

### Implications of nephron-to-stromal lineage crosstalk in normal development and disease

The nephrogenic niche consists of a specialized microenvironment that has been shown to regulate the balance of progenitor cell maintenance/differentiation in coordinating nephrogenesis. The *Foxd1*+ stromal progenitor population localized to this niche is known to play a critical role in establishing this microenvironment and gives rise to the diverse population of interstitial cells necessary for normal kidney development (Wilson & Little, 2021). Here, we show that defects in nephrogenesis, including those caused by mutations in *Wt1*, also result in abnormal stromal development. This is of particular interest given that loss-of-function of WT1 is associated with a number of renal pathologies, including Denys Drash syndrome, WAGR (Wilms tumor, aniridia, genitourinary anomalies, and a range of developmental delays) syndrome, and predisposition to Wilms tumor (Torban & Goodyer, 2024). While Wilms tumors with *WT1* mutations have been associated with stromal predominant histology (Hastie, 2017), previous investigations of *Wt1* mutations targeted to the embryonic renal stroma have not supported this mechanism in Wilms tumorigenesis (Huang et al., 2016; Weiss et al., 2020). However, in this study, we specifically examined *Wt1* mutation targeted to the NPC lineage and interestingly identify non-autonomous effects resulting in an abnormal expansion of the stromal progenitor population. Given that the nephrogenic stroma normally signals to the NPCs, its abnormal expansion may potentially cause aberrant crosstalk within the nephrogenic niche, which may be of interest to explore in future studies. Furthermore, understanding the mechanisms of how such cell-lineage crosstalk coordinates progenitor cell maintenance/differentiation may lead to a better understanding of how the stroma contributes to generating “higher-order” structures in kidney organoids to guide engineering strategies recapitulating in-vivo nephrogenesis.

While our analyses focused on signaling between the nephron and stromal lineages, abnormal signaling from the UB may be another possible mechanism affecting the disrupted stromal development in *Six2cre;Wt1^c/c^, Wnt4-null,* and *Six2cre;RosaDTA^c/+^* mutant kidneys given that all three models show decreased UB branching and global defects in development that may be reflective of an “immature, earlier stage”. To partially address such effects of decreased UB branching and a “delayed” developmental phenotype, we evaluated the expression of *Foxd1*+ stromal markers at E12.5 and E13.5 as described above (Fig. 3 and Supplemental Fig. S2). These findings suggest that the abnormal expansion of *Foxd1* and *Ntn1* expression in mutant kidneys is not simply due to decreased UB branching, as it is not seen in E12.5 or E13.5 kidneys. However, it is still possible that signaling from abnormally developed UBs (either from the inability to branch normally or from altered development due to disrupted signaling from the NPC lineage) may contribute in the disruption of stromal development in the mutant kidneys. Nonetheless, the findings from this study highlight the coordination of progenitor cell maintenance/differentiation in the developing kidney and provide new insights into the heterogeneity and regulation of the *Foxd1*+ stromal progenitor population.

## MATERIALS AND METHODS

### Animal models

All animals were housed, maintained, and used according to National Institutes of Health (NIH) and Institutional Animal Care and Use Committees (IACUC) approved protocols at the University of Texas Southwestern Medical Center (OLAW Assurance Number D16-00296). Mouse lines used in this study include: *Six2creTGC*, Jax strain #009606 (Kobayashi et al., 2008), *Wt1^c/c^*, Jax strain #019554 (Gao et al., 2006), *DTA^c/c^*, Jax strain #006331 (Brockschnieder et al., 2004), and *Wnt4-null*, Jax strain #002866 (Stark et al., 1994). All mice were bred on a mixed genetic background. For experimental assays, females homozygous for conditional alleles were crossed with male cre-line mice, with the day of plug counted as E0.5. Pregnant females were sacrificed at various gestational time points. Lineage-tracing experiments were performed by crossing *Rosa26^EYFP^,* Jax strain #006148 (Srinivas et al., 2001) reporter mice with the above mouse lines. Mice with the desired genotype were randomly selected regardless of sex. Cre-negative littermates were used as controls.

### Kidney sample preparation, histology, immunofluorescence (IF), and in-situ hybridization (ISH)

Embryonic tissue was fixed in 4% paraformaldehyde, embedded in paraffin, sectioned into 5 μm slices, and subjected to Hematoxylin and Eosin staining or IF. Slides for IF were immersed and boiled with either 10 mM sodium citrate or TE antigen retrieval buffer and blocked with a solution of 5% normal donkey serum for 1 hour at room temperature followed by the application of primary antibodies diluted in blocking solution. The following primary antibodies were used: SIX2 (Proteintech; 11562-1-AP; 1:200 dilution, Rabbit), LXH1 (DHSB; 4F2; 1:100 dilution; Mouse), NCAM (Sigma; C9672; 1:200 dilution; Mouse), CK (DSHB; Troma-III; 1:50 dilution; Rat), MEIS 1/2/3 (Active Motif; 39795; 1:100 dilution; Mouse); ALDH1A2 (Sigma; HPA010022; 1:200 dilution; Rabbit), DBA (Vector; B-1035; 1:500 dilution; Biotinylated), and GFP (Aves; GFP-1020; 1:200 dilution; Chicken). Immunofluorescence staining of control and mutant paraffin sections was carried out on the same slide, and visualization was carried out with the same microscope/photography settings (NikonA1 inverted confocal microscope). For in-situ hybridization assays, tissue was fixed with 4% paraformaldehyde, cryoprotected with 30% sucrose, embedded in OCTmedium (TissueTek), sectioned into 10 μm sections, rehydrated with PBS before being treated with 15 μg/ml proteinase K for 10 min and fixed in 4% PFA followed by an acetylation step. Slides were then washed and incubated with pre-hybridization buffer for 1 hour at room temperature before being hybridized with the specific probe overnight at 65°C. Slides were then washed in 0.8X SSC then transferred to NTT before blocking with 2% blocking solution (Roche) for at least 1 hour at room temperature. Slides were then incubated with anti-Dig alkaline phosphatase-conjugated antibody (Roche, 1:4000) overnight at 4°C. The next day, slides were washed in three times in NTT and three times in NTTML before incubating with BM purple (Roche); slides were then fixed with 4% PFA and mounted using glycergel (Dako).

### Single nuclei RNA-sequencing

Nuclei were isolated as previously reported (Wu et al., 2018) from paired kidneys harvested from E15.5 embryos with Nuclei EZ Lysis buffer (Sigma NUC-101) supplemented with protease inhibitor (Roche 5892791001) and RNase inhibitor (Promega N2615, Life Technologies AM2696). Samples were homogenized using a Dounce homogenizer (Kimble Chase 885302-0002) in 2ml of ice-cold Nuclei EZ Lysis buffer first with a loose pestle, then passed through a 200 µm strainer into 50 ml conical tube and followed by homogenization with a tight pestle which was then filtered through a 40-µm cell strainer (pluriSelect 43-50040-51) and incubated on ice for 5 minutes. The suspension was centrifuged at 500 g x for 5 min at 4 °C with the pellet resuspended and washed with 4 ml of the buffer and incubated on ice for 5 min. After another centrifugation, the pellet was resuspended with Nuclei Suspension Buffer (1x PBS, 0.07% BSA, 0.1% RNase inhibitor) and filtered through a 5-µm cell strainer (pluriSelect 43-50020-50) into a conical tube. The nuclei were counted with a disposable hemacytometer and diluted to a concentration of 700-1200 cells per µL. Nuclei were sequenced using the 10x Chromium Single Cell Platform (10x Genomics) targeting 5,000 nuclei per sample and 50,000 reads per nuclei. Library prep and sequencing was completed by the UT Southwestern McDermott Sequencing core using the 3’ GEX kit on Next Seq (5-150 cycle high output) and aligned to GRCm39.

### Bioinformatic analyses of single nuclei RNA-sequencing

The 10X Genomics data were analyzed using the Cell Ranger Pipeline cellranger (Version 5.0.1) for alignment, quantification, and initial preprocessing of single nuclei RNA-seq data. The reads were aligned using STAR to the mouse reference genome provided by 10XGenomics (refdata-gex-mm10-2020-A). The estimated cell number was derived by plotting the UMI counts against the barcodes. Further filtering of the expression data matrices was done to ensure high quality data. Cells with a minimum number of UMI >= 500, a minimum number of detected genes >= 250, log10GenesPerUMI > 0.80 and mitoRatio < 0.20, were selected. Cells were clustered using the Louvain algorithm implemented in Seurat based on the top principal components (Butler et al., 2018; Hao et al., 2021; Satija et al., 2015; Stuart et al., 2019). Both Seurat (4.0.5) and CZ CellxGene (1.3.1) were used for clustering analysis, differential expression analysis, and visualization of single-cell RNA-seq data (https://doi.org/10.1101/2021.04.05.438318; https://www.biorxiv.org/content/10.1101/2020.08.28.270652v2). Cell clusters were identified based on marker gene expression and visual inspection (Chaney et al., 2022). Publicly available human fetal kidney snRNAseq data was re-processed using Seurat (version 4.3.0). Raw counts were log-transformed, and the top 4000 variable genes were selected for scaling, principal component analysis, and batch correction with Harmony (version 1.2.0). The top 20 dimensions were selected for downstream processing, including UMAP and Louvain clustering. Clusters were calculated at multiple resolutions and an optimal resolution was selected based on examination of a clustree (version 0.5.1) plot and marker gene lists.

### Statistical analysis and data visualization

Statistics for the bioinformatic analyses of the single nuclei RNA-seq data included: statistical significance of differentially expressed genes (DEG) lists of control and mutant determined by Wilcoxon Rank-Sum Test with significance thresholds adjusted for multiple testing (< 0.05) with the data filtering/analyses as described above. Data presented in the figures are representative images from one of at least three different experiments on different embryos/organs. No significant variability was noted in tissues of the same genotype; all animals with correct genotypes were included in the analysis. ImageJ was used to measure nephrogenic stromal width on immunofluorescence images stained with ALDH1A2 by measuring the distance from the outer cap mesenchyme (i.e., SIX2 posivite cells) to the periphery of the kidney on sections of control and mutant kidneys, with 10 measurements obtained from 3 separate embryos of each sample, and statistical significance evaluated by one way ANOVA and Tukey HSD used for group comparison.

## ACKNOWLEDGEMENTS

This research was supported in part by the computational resources provided by the BioHPC supercomputing facility located in the Lyda Hill Department of Bioinformatics, UT Southwestern Medical Center, TX. URL: https://portal.biohpc.swmed.edu. We would also like to acknowledge Dr. Thomas Carroll for providing the *Foxd1* in-situ probe and mouse lines used in this study.

## DATA AVAILABILITY

All data, raw and processed files, are available upon request. Single nuclei RNA-seq from mouse control kidneys will be made available in the Gene Expression Omnibus (GEO) data repository prior to publication. Human fetal kidney snRNA-seq is available via GEO (GSE232479) as previously published (Kim et al., 2024).

## SUPPLEMENTAL MATERIAL LIST

**Supplemental Figure S1.**
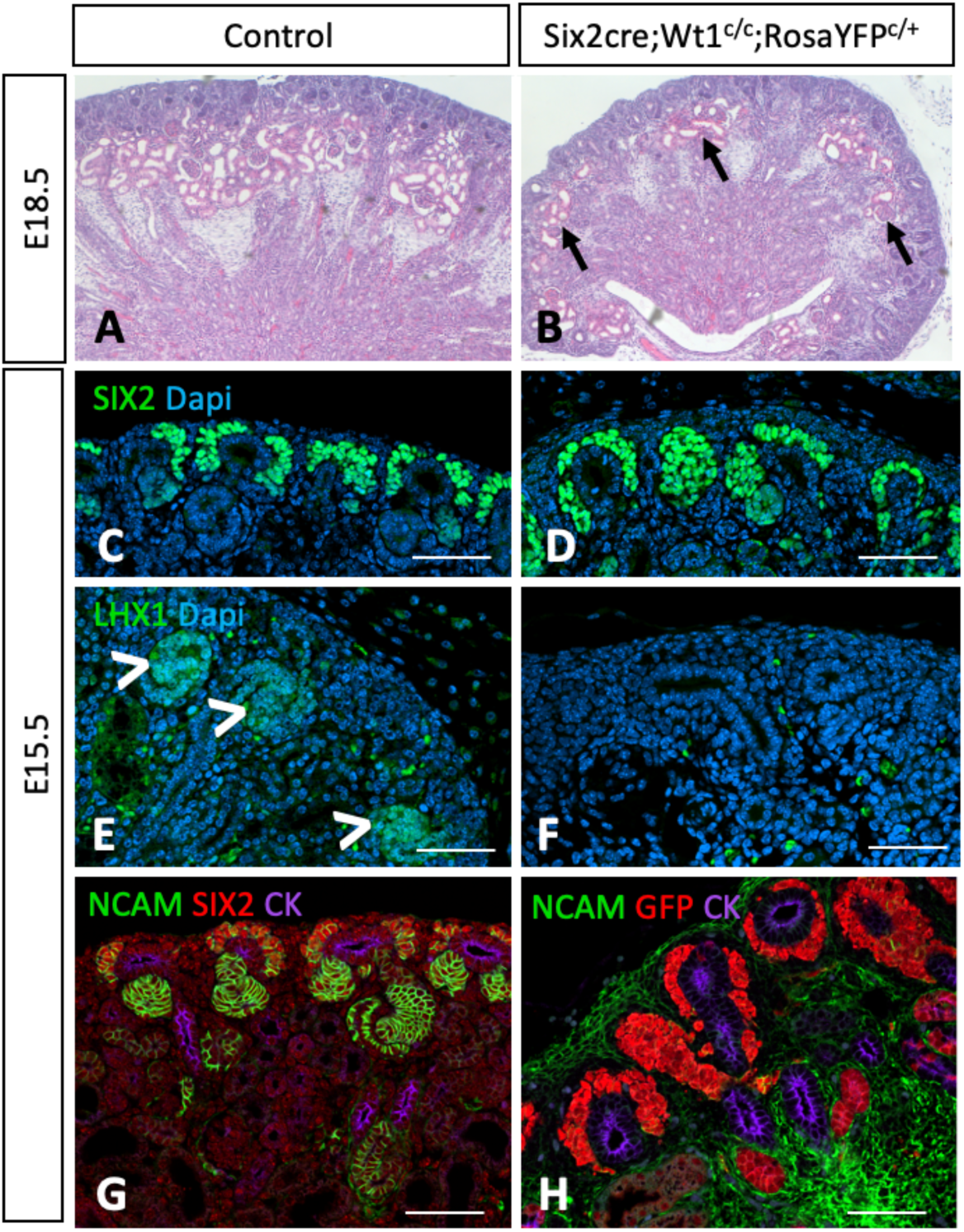
Characterization of the *Six2cre;Wt1^c/c^* mutant mouse model. (A-B) Histology of E18.5 cre-negative control kidneys (A) is shown in comparison to *Six2cre;Wt1^c/c^* mutant kidneys with defects in nephrogenesis, though some proximal tubules and glomeruli are observed (B, arrows) presumably due to incomplete efficiency of the *Six2cre* model. (C-H) Immunofluorescence of E15.5 control and mutant kidneys show NPCS (labed with SIX2) maintained in mutant kidneys (D) with early differentiating nephron structures (labeled with LHX1, arrowheads) lost in mutant kidneys (F). Inclusion of a *Rosa26^EYFP^* reporter (which labels with GFP antibody) confirms that *Six2cre* specifically targets the NPC lineage with no recombination observed in the stroma as expected (H).

**Supplemental Figure S2.**
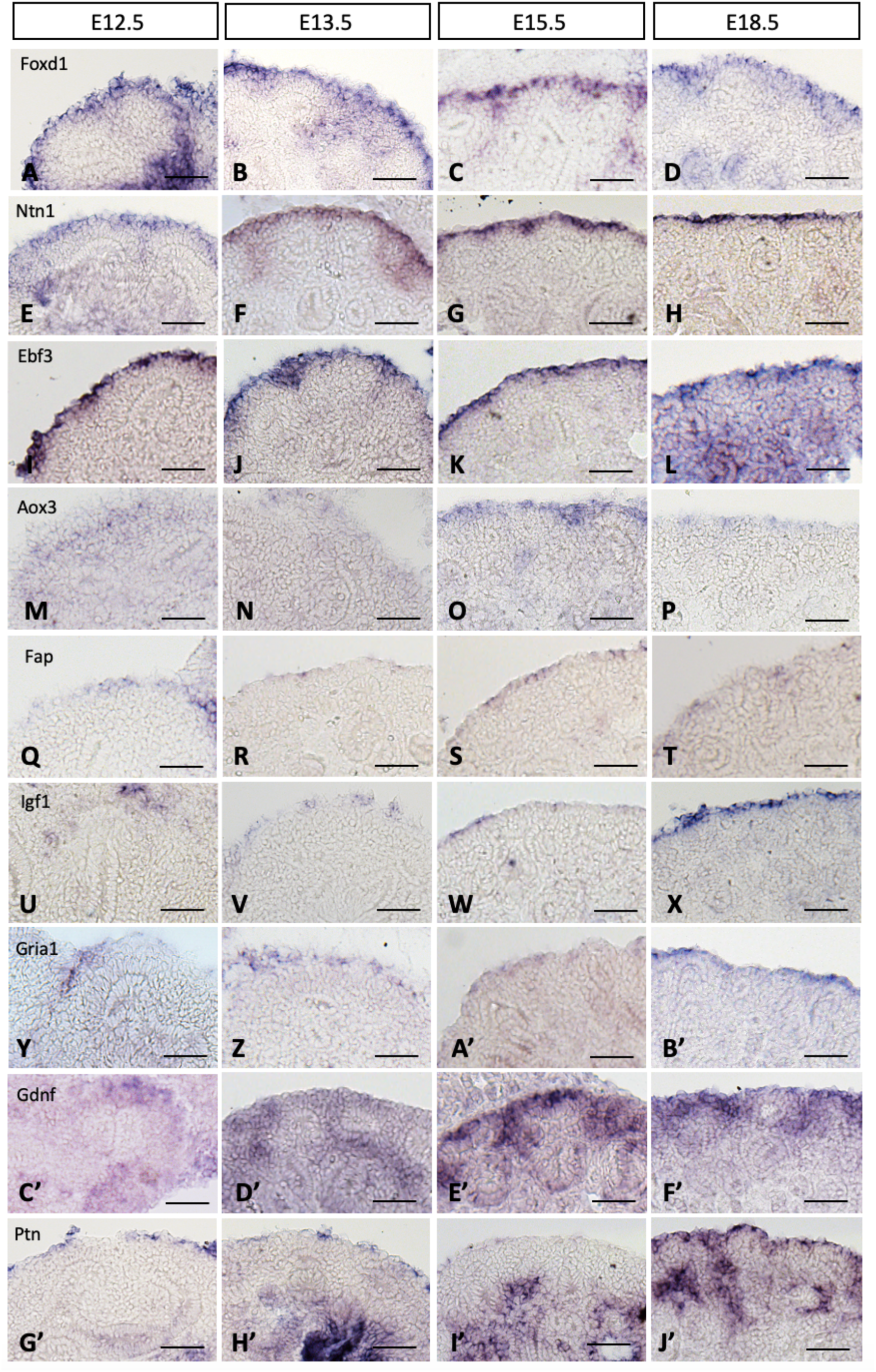
Expression of stromal progenitor cell markers throughout time points of normal kidney development. Identified markers of the *Foxd1*+ stromal progenitor population from snRNA-seq were evaluated at E12.5, E13.5, E15.5, and E18.5 in control kidneys.

**Supplemental Figure S3.**
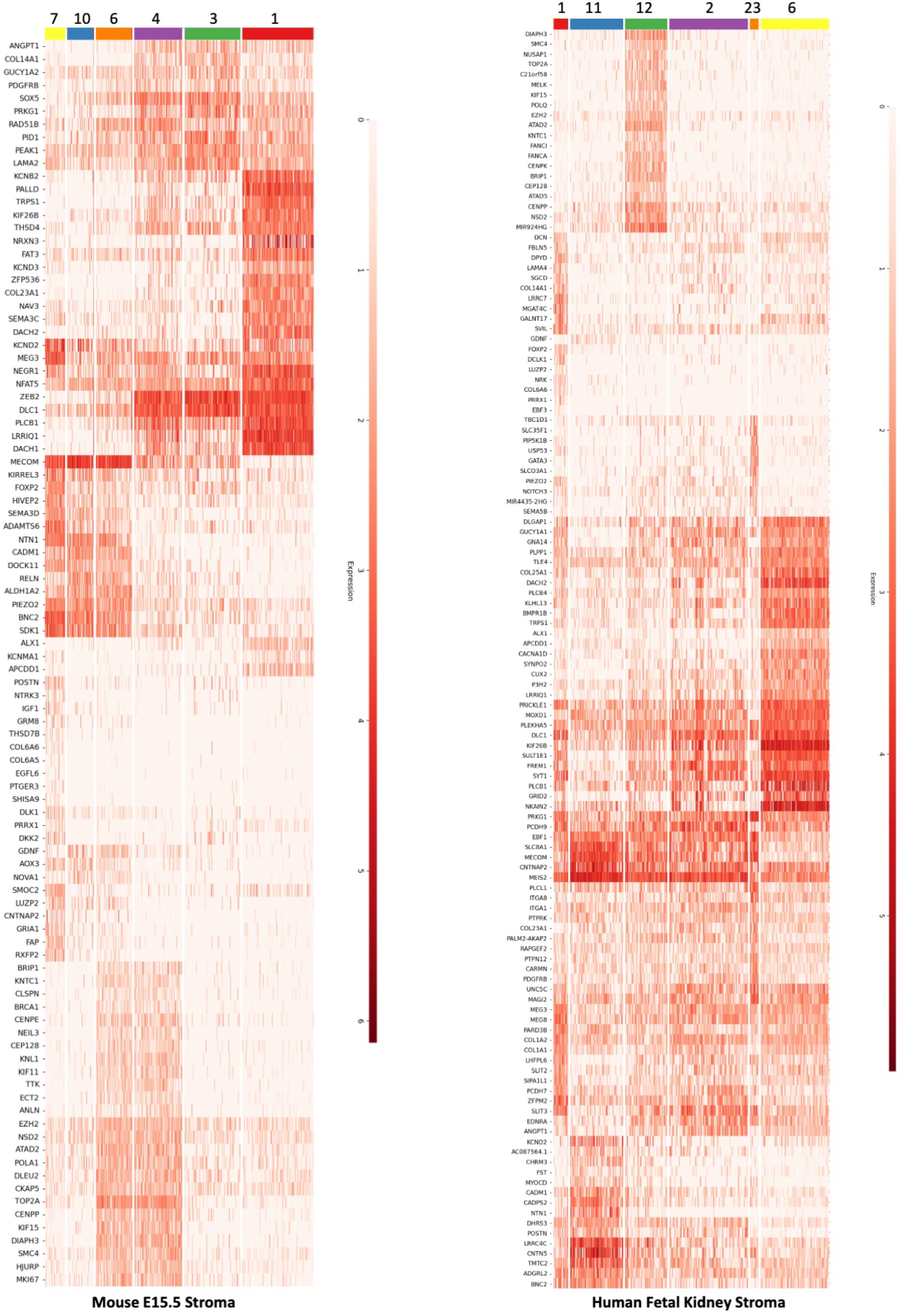
Heatmap of stromal clusters identified in murine and human fetal kidneys. Analysis of snRNA-seq of murine and human fetal kidneys identified six subclusters of stroma, with transcriptional profiles of marker genes and top differentially expressed genes shown via heatmap.

**Supplemental File S1.** E15.5 mouse snRNA-seq_Marker genes.xlsx

**Supplemental File S2.** E15.5 mouse snRNA-seq_DEG for Cre-neg vs Six2cre_combined nephrogenic zone stromal clusters 7, 10, and 6.xlsx

**Supplemental File S3.** E15.5 mouse snRNA-seq_DEG for clusters 7, 10, and 6.xlsx

**Supplemental File S4.** Human fetal kidney snRNA-seq_Marker genes.xlsx

